# Neural timescales reflect behavioral demands in freely moving rhesus macaques

**DOI:** 10.1101/2023.03.27.534470

**Authors:** Ana M.G. Manea, Anna Zilverstand, Benjamin Hayden, Jan Zimmermann

**Author notes:** Corresponding author: Ana Manea Department of Neuroscience, University of Minnesota Minneapolis, MN, 55455.

## Abstract

Previous work has demonstrated remarkably reproducible and consistent hierarchies of neural timescales across cortical areas at rest. The question arises how such stable hierarchies give rise to adaptive behavior that requires flexible adjustment of temporal coding and integration demands. Potentially, this previously found lack of variability in the hierarchical organization of neural timescales could be a reflection of the structure of the laboratory contexts in which they were measured. Indeed, computational work demonstrates the existence of multiple temporal hierarchies within the same anatomical network when the input structure is altered. We posit that unconstrained behavioral environments where relatively little temporal demands are imposed from the experimenter are an ideal test bed to address the question of whether the hierarchical organization and the magnitude of neural timescales reflect ongoing behavioral demands. To tackle this question, we measured timescales of local field potential activity while rhesus macaques were foraging freely in a large open space. We find a hierarchy of neural timescales that is unique to this foraging environment. Importantly, although the magnitude of neural timescales generally expanded with task engagement, the brain areas’ relative position in the hierarchy was stable across the recording sessions. Notably, the magnitude of neural timescales monotonically expanded with task engagement across a relatively long temporal scale spanning the duration of the recording session. Over shorter temporal scales, the magnitude of neural timescales changed dynamically around foraging events. Moreover, the change in the magnitude of neural timescales contained functionally relevant information, differentiating between seemingly similar events in terms of motor demands and associated reward. That is, the patterns of change were associated with the cognitive and behavioral meaning of these events. Finally, we demonstrated that brain areas were differentially affected by these behavioral demands - i.e., the expansion of neural timescales was not the same across all areas. Together, these results demonstrate that the observed hierarchy of neural timescales is context-dependent and that changes in the magnitude of neural timescales are closely related to overall task engagement and behavioral demands.

## Introduction

Behavioral coordination and adaptation across an ever-changing environment are a hallmark of cognition in biological systems. To function in our daily lives we simultaneously consider auditory, visual, and sensory input while achieving motor coordination, each of which spans a continuum of spatial and temporal scales. Consider one of the most studied systems in neuroscience, the visual cortex. It is well known that neurons along the visual pathway have increasingly larger receptive fields (Hubel & Wiesel, 1968) —higher-level visual areas respond to information from large portions of space by integrating input from neurons in the early visual cortex which possess smaller receptive fields. Events not only unfold over multiple spatial but also over a multitude of temporal scales (Kohn, 2007). In fact, spatial integration in the visual system would not be possible without temporal integration of information, and implicitly, neurons simultaneously processing input across multiple timescales (Cocchi et al., 2016; Zhou et al., 2018). Indeed, this hierarchical increase in representational complexity is closely followed by a hierarchy of longer temporal processing windows (Ogawa & Komatsu, 2010). Similarly, this hierarchy of neural processing across different scales of complexity is found in the prefrontal cortex (PFC), with progressively more abstract representations and higher-order control on a posterior-anterior and ventral-medial axes (Badre, 2008; Fuster, 2001; Koechlin et al., 2003). Moreover, electrophysiological, and functional Magnetic Resonance Imaging (fMRI) results in human and nonhuman primates at rest have demonstrated that the frontal lobe is organized along a hierarchical gradient of neural timescales that mirrors its functional architecture (Cavanagh et al., 2016; Maisson et al., 2021; Manea et al., 2022; Murray et al., 2014; Raut et al., 2020). These parallel results suggest that neural timescales in the PFC might be functionally relevant, nevertheless, direct evidence to support this conclusion is limited.

Generally, it is thought that areas operating at slower timescales have a wider temporal processing window to integrate information from other brain areas (Cocchi et al., 2016; Hasson et al., 2008; Honey et al., 2012; Ito et al., 2020; Soltani et al., 2021). Indeed, prefrontal brain areas with more abstract representations that display relatively more information integration compared to other brain areas generally also display slower neural timescales (Maisson et al., 2021; Manea et al., 2022). Based on these findings, it could be argued that neural timescales merely reflect a relatively static property inherited from their place in the anatomical hierarchy that allows neurons within that area to integrate over a stable temporal scale. In this case, although neural timescales would facilitate function, the timescales themselves would not change in a functional manner. Alternatively, it could be argued that anatomy instead might impose a range of timescales that bounds the dynamics over which brain areas can operate. For example, it is conceivable that there are stable hierarchies of timescales that can expand and contract depending on functional demands.

Findings on the dynamics of neural timescales in the context of behavior are scarce (see Cavanagh et al., 2020; Gao et al., 2020; Zeraati et al., 2021). There is currently conflicting evidence about the behavioral dependence of neural timescales, their hierarchical organization and general function. Some preliminary evidence suggests that neural timescales expand with task engagement and attention, and hence are potentially functionally relevant (Gao et al., 2020; Zeraati et al., 2021). In contrast, a plethora of previous findings have demonstrated remarkably reproducible and consistent neural timescales across cortical areas at rest (see **Fig. 1A;** Cavanagh et al., 2016, 2018; Cirillo et al., 2018; Fascianelli et al., 2019; Maisson et al., 2021; Murray et al., 2014; Nougaret et al., 2021; Wasmuht et al., 2018), with one study even concluding that the hierarchy of neural timescales appears invariant to task context and that neural timescales are not affected by behavioral demands (Rossi-Pool et al., 2021). The question that arises from these findings is how such apparently static temporal properties can accommodate adaptive and flexible behavior. After all, not all behavior follows the same exact temporal sequence with the same temporal scale. Indeed, a large-scale dynamical model of the macaque neocortex exhibits not one, but multiple temporal hierarchies, as indicated by unique responses to visual and somatosensory stimulation (Chaudhuri et al., 2015). The existence of multiple concurrent neural timescales gradients that are dynamically expressed (Chaudhuri et al., 2015; Li & Wang, 2022; Spitmaan et al., 2020) might support behavioral changes at different temporal scales. This would not be surprising since even sheer neuronal variability is ubiquitous and functional not only across but also within brain areas, and it is well established that neural processing displays complex temporal dynamics (Churchland et al., 2010; Goris et al., 2014, 2015). Indeed, although characteristic timescales have been assigned to brain areas as a whole, single neurons display heterogeneous neural timescales at rest (Cavanagh et al., 2016; Cirillo et al., 2018; Fascianelli et al., 2019; Fontanier et al., 2022; Murray et al., 2014; Wasmuht et al., 2018). The question arises whether this heterogeneity is purely anatomical or whether it is the result of both anatomy and contextual demands (Runyan et al., 2017). Together, while the majority of findings on neural timescales at rest have found a similar cortical hierarchy across studies, computational work and recent empirical findings suggest that neural timescales and their hierarchical organization may be context-dependent, varying with behavioral and environmental demands.

**Fig. 1.**
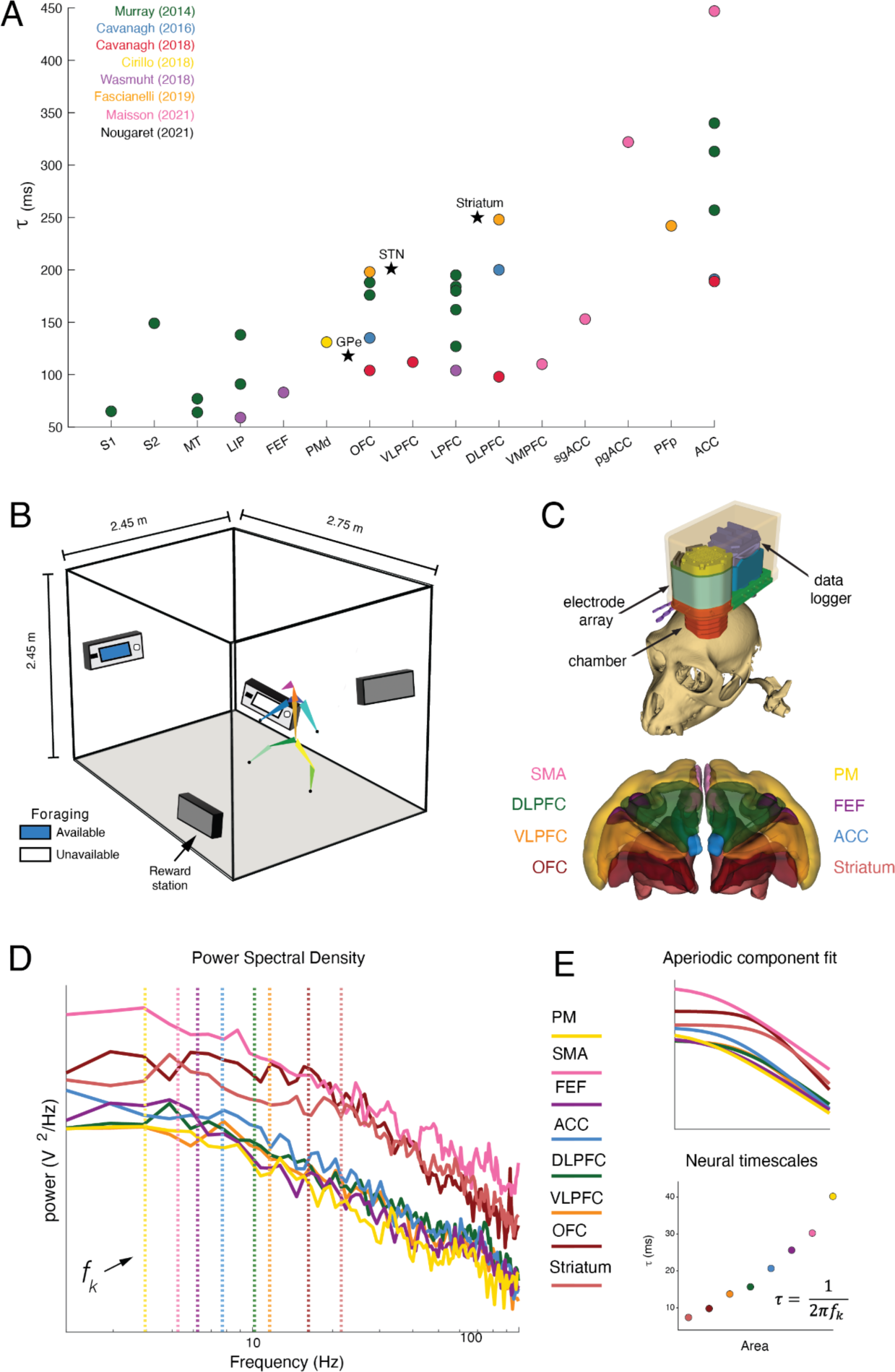
Overview of neural timescales and experimental design. **(A)** Hierarchical organization of neural timescales at rest (τ) estimated from neuronal spiking data. Neural timescales were estimated in 14 cortical and 3 subcortical areas). Traditionally, neural timescales were estimated in the pre-trial period of various tasks (i.e., chaired electrophysiology) by fitting an exponential decay function to the autocorrelation function (i.e., time lagged correlation). Each circle represents the population-level τ for each cortical area and the stars represent population-level τ for each subcortical area reported in each study. **(B)** Depiction of the cage and foraging task. The subjects were allowed to freely explore and interact with reward stations in an open space - i.e., 2.45 x 2.45 x 2.75 m cage with barrels. **(C)** Our recording system and recording sites: dorsal striatum, OFC, VLPFC, DLPFC, ACC, FEF, PM, SMA **(D)** Local field potential (LFP) power spectral densities (PSDs) from example channels in Subject W. **(E) Top**: Aperiodic component fit for the example PSDs. We applied spectral parameterization to infer timescales from the PSDs (Donoghue et al., 2020). The periodic oscillatory peaks were discarded and the “knee frequency” (*f_k_* vertical dashed lines) was extracted from the fit of the aperiodic component. **Bottom:** Neural timescales (τ) were inferred from *f_k_* via the embedded equation.

We propose that the failure to find multiple hierarchies of neural timescales in previous experiments may be a by-product of the rigid structure of traditional experiments that enforce stationary temporal scales. In particular, investigating neural timescales in a constrained lab environment (i.e., chaired electrophysiology) with trialized tasks, imposes bounded temporal structure and limits the complexity of the input entering and the output leaving the brain. If these constraints were removed, neural and behavioral dynamics would become temporally unconstrained —except for the boundaries imposed by biophysics. Therefore, what we know about neural timescales might not be entirely intrinsic to the biological system, but rather a reflection of the conditions imposed by the structure of the experimental paradigm (Hastings, 2010; Marom, 2010). While this approach has brought invaluable contributions to our understanding of the brain and behavior, it is nonetheless limited when it comes to studying the functional relevance of neural timescales. To understand and establish a neurobehavioral timescale correspondence, it is imperative to approach the question from a more unconstrained perspective. Here, we thus investigated neural timescales in an unconstrained experimental paradigm with relatively minimal temporal structure.

We hypothesized that the observed hierarchy of neural timescales is dependent on the environment, as reflected by the temporal constraints, the complexity and nature of the input, and required motor output. Moreover, we hypothesized behaviorally-dependent shifts in the magnitude of neural timescales —and hence, dynamic coordination between neural timescales and the temporal scales over which behavior unfolds. To test these hypotheses, we investigated how the brain handles multi-scale signals to drive purposeful behavior while rhesus macaques were performing a foraging task. We were able to use detailed three-dimensional behavioral tracking while the monkeys were free to move and forage in a large open field environment (Bala et al., 2020). Our experimental paradigm imposed minimal temporal constraints and put emphasis on self-paced behavior rather than focusing on a particular cognitive or perceptual process in isolation. We simultaneously recorded brain activity in eight brain areas: orbitofrontal cortex (OFC), ventrolateral prefrontal cortex (VLPFC), dorsolateral prefrontal cortex (DLPFC), frontal eye fields (FEF), anterior cingulate cortex (ACC), premotor (PM) cortex, supplementary motor area (SMA), and dorsal striatum. We investigated neural timescales from a population perspective as reflected in the local field potential (LFP) activity —in particular, we estimated the aperiodic timescales which correspond to the exponential decay timescales measured in previous studies (Gao et al., 2020; Halgren et al., 2021). In contrast to previous studies investigating neural timescales, our environment not only involved the integration of highly complex multi-modal input but also required complex motor output. We found a hierarchy of neural timescales unique to this foraging environment. Next, we demonstrated that neural timescales expand with task engagement although the areas’ relative position in the hierarchy remains the same across the recording session. Finally, we showed that the change in neural timescales is dynamic and reflects the abstract meaning of foraging events. Together, this demonstrates that while anatomy constrains the space of possible neural timescales, the observed hierarchical organization and magnitude of neural timescales is heavily dependent on behavioral demands.

## Results

Two macaques performed a foraging task in a large open space that allowed for unconstrained movement (**Fig. 1B** and **Methods**). The environment contained four reward stations positioned at fixed locations. The reward stations dispensed 1.5 mL of liquid reward for each of the first four lever presses and became unavailable for 3 minutes after the fifth lever press (i.e., depleted reward station; see **Methods** for task details). The average daily recording session was 97.8 minutes (SD ± 5.2 minutes).

We tracked the position of thirteen joints (key points) in our subjects with OpenMonkeyStudio (Bala et al., 2020). We recorded neural activity using a data logger (SpikeGadgets, San Francisco, USA) attached to a multi-electrode array (Gray Matter Research, Bozeman, USA) with 128 independently movable electrodes. We recorded both isolated neurons and LFPs from eight areas: orbitofrontal (OFC), ventrolateral prefrontal (PFC), anterior cingulate (ACC), dorsolateral prefrontal (DLPFC) cortices, frontal eye fields (FEF), supplementary motor area (SMA), premotor cortex (PM) and the dorsal striatum (**Fig. 1C**). To quantify neural timescales, we focused on the LFPs because of their spatial coverage (see **Fig. 1D** and **Methods**).

### Neural timescales are variable

We demonstrated that the hierarchy of neural timescales is (1) is different from previous findings, and hence dependent on the experimental paradigm used and (2) stable within our environmental context across all sessions. We further found that (3) while areas in our unconstrained foraging task exhibited the same relative position in the neural timescales’ hierarchy throughout our recording sessions, the magnitude of the neural timescales changed depending on task demand. We demonstrated that the non-stationarity of neural timescales was related to task engagement, and to a lesser extent to speed of movement–more engagement was accompanied by slower temporal dynamics in all areas.

First, we examined session-wide neural timescales for individual recording sessions. For each recording site, we estimated neural timescales using a 10 sec moving window with 5 sec overlap (see **Fig. 2A** and **Methods**). To identify the neural timescales characteristic of each brain area, we computed the median neural timescales collapsing across sessions and subjects. We demonstrated that neural timescales estimated from LFPs are ∼10 times faster than those estimated from neuronal spiking data — i.e., ∼10-50 ms, which is consistent with previous findings (Gao et al., 2020). We further found that the magnitude of neural timescales decreased systematically across the duration of the recording session in all areas (p < 0.05, linear regression model, **Fig. 2A**).

**Fig. 2.**
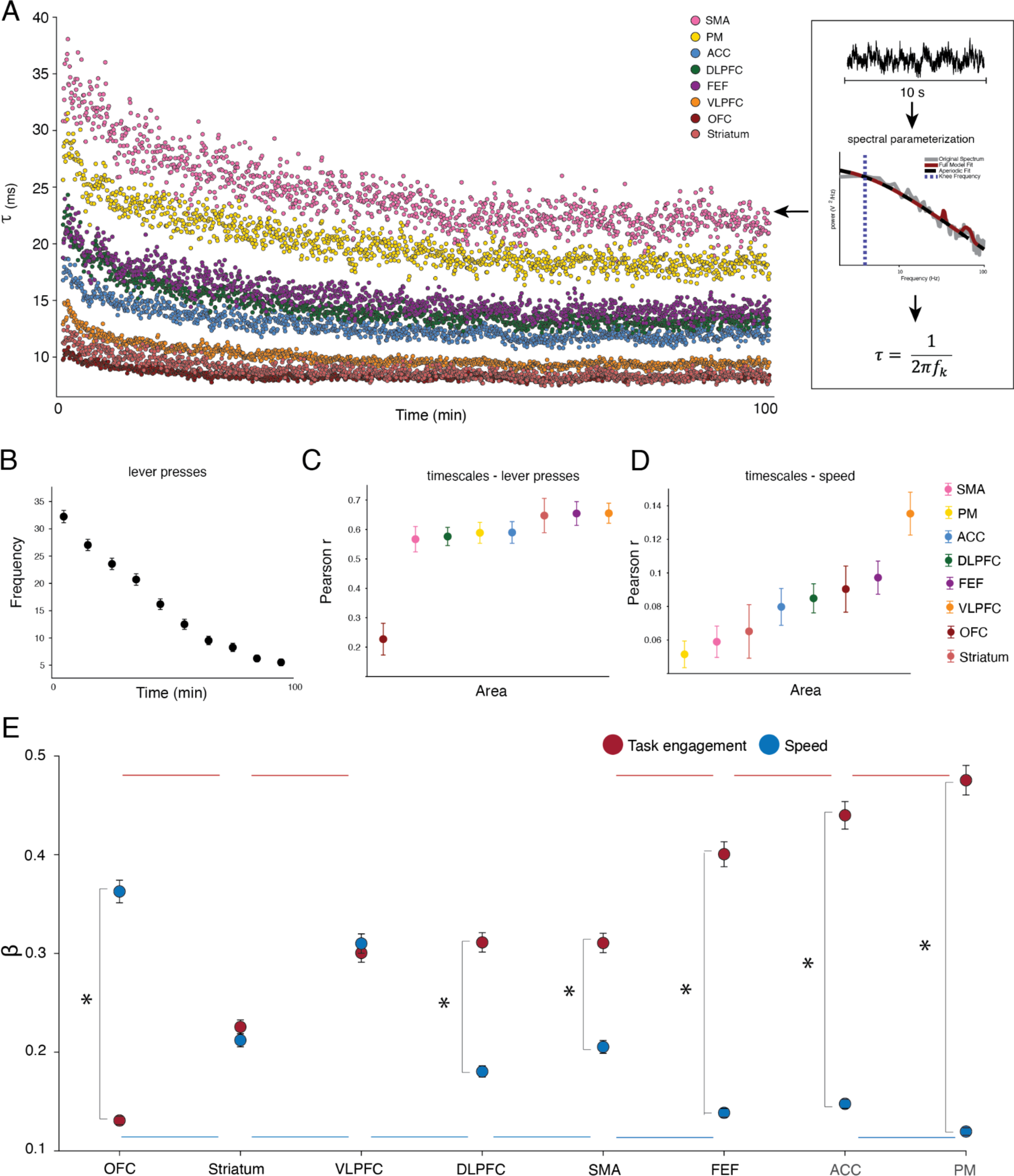
Session-wide timescale dynamics and correspondence to behavioral variables. **(A) Right:** Neural timescales (τ) estimated for individual recording sessions from Power Spectral Densities (PSDs) **Left**: Neural timescales across time, collapsed across recording sessions and subjects. The hierarchical ordering of the areas is conserved across the duration of the recording session. **(B)** Frequency of lever pressing, a putative index of task engagement, decreases over time. Y-axis: average number of lever presses in each time bin. Bars indicate standard error of the mean across recording sessions. **(C)** The number of lever presses, a putative index of task engagement, is correlated with neural timescales in all areas. Y-axis: correlation coefficient between number of lever presses and neural timescales across time. Circles: brain areas, median ± s.e.m across recording sessions. **(D)** Speed of movement and neural timescales are weakly correlated. Circles represent brain areas, median ± s.e.m across sessions. **(E)** The effect of task engagement is gradually stronger and more separable from the effect of speed as we move from ventral to dorsal areas. Y-axis: average regression coefficients. Circles represent brain areas, mean ± s.e.m across iterations. Horizontal lines: Statistical significance at p < 0.05 between brain areas (Red: task engagement; Blue: speed). Asterisk: statistical significance between predictors within an area at p < 0.05.

We observed that task engagement (i.e., lever presses) also decreased across the recording session in both animals (**Fig. 2B**). Hence, we asked whether neural timescales and task engagement were related across the recording session. We hypothesized that there is a monotonic relationship between the two, with more task engagement being accompanied by slower dynamics. Specifically, we divided each session into ten equally sized (∼10 minute-long) segments. As a proxy for task engagement, we calculated the total number of lever presses in each bin. For every recording session, we computed the correlation between the magnitude of the neural timescales and our task engagement index. For all areas, we observed a strong positive relationship between task engagement and the magnitude of the neural timescales - the median Pearson correlation coefficient across sessions ranging between 0.23 in the OFC and 0.65 in the VLPFC (**Fig. 2C**). To assess whether the median of the obtained distribution of Pearson correlation coefficients was significantly larger than 0, we performed a one-sample Wilcoxon signed rank for every area. In all areas, the median Pearson correlation coefficient was significantly larger than 0 (p < 0.05, with Bonferroni correction).

Next, we asked whether neural timescales were related to movement, to account for the possibility that changes in neural timescales could be driven by increased motor activity, during intervals of high displacement or motion in general. For every recording session, we computed the correlation between speed of displacement and neural timescales. We found that neural timescales were only weakly correlated with movement speed - the median Pearson correlation coefficient across sessions ranging from 0.05 in the PM cortex to 0.14 in the VLPFC (**Fig. 2D**). To assess whether the median of the obtained distribution of Pearson correlation coefficients was significantly larger than 0, we performed a one-sample Wilcoxon signed rank for every area. In all areas, the median Pearson correlation coefficient was significantly larger than 0 (p < 0.05, with Bonferroni correction). In combination, these results suggest that the gradual decrease in neural timescales throughout the recording session was primarily related to task engagement, an aggregate of behavioral state parameters in our task, and less so to movement parameters such as speed, a hypothesis we will further test next.

To test the unique relationship between speed, task engagement and neural timescales for the brain areas we recorded from, as well as to control for temporal autocorrelations, we next performed the following analysis. For each area, we randomly sampled without replacement n (i.e., equivalent to the number of sessions) observations out of the total number of data points (note: the total number of data points per area can be calculated as the number of sessions x number of bins). For each subsample, we fit a linear regression model (see **Fig. 2 Supplement 1** for the resulting distributions of standardized β coefficients). We demonstrated that the effect of task engagement was gradually stronger in more dorsal areas - i.e., the regression coefficients are progressively larger (p < 0.05, pairwise independent sample t-test; **Fig. 2E**) displaying the following ventro-dorsal ordering: OFC < Striatum < VLPFC-DLPFC-SMA < FEF < ACC < PM. Conversely, the effect of speed was gradually stronger in ventral areas - i.e., the regression coefficients are progressively larger (p < 0.05, pairwise independent sample t-test; **Fig. 2E**) displaying the following dorso-ventral ordering: OFC > VLPFC > Striatum - SMA > DLPFC > FEF - ACC > PM. Moreover, the effect of task engagement was larger than that of speed in more dorsal areas: SMA, PM, FEF, ACC, DLPFC (p < 0.05, two-sided paired t-test with Bonferroni correction). In contrast, there was no significant difference between the two predictors in VLPFC and the striatum and the relationship was reversed in the OFC (p < 0.05, two-sided paired t-test with Bonferroni correction).

In summary, we found a ventro-dorsal hierarchy that partially overlapped but also deviated from previous findings in important ways. We demonstrated a monotonic relationship between brain areas and session-wide neural timescales (b = 2.29, 95% CI = [2.27 2.31]; monotonic Bayesian regression model) with the following hierarchical organization: OFC < Striatum < VLPFC < ACC < DLPFC < FEF < PM < SMA (**Fig. 2A**). Our observed neural timescales were faster in ventral areas and slower in more dorsal areas. To systematically assess the stability of this hierarchy across time, we additionally ranked the areas at each time point and compared their ordering to the session-wide hierarchy of neural timescales by using Spearman rank correlation (average Spearman rank correlation coefficient 0.99, SD 0.01). This analysis further supported the Bayesian regression results, demonstrating a monotonic relationship between areas, with the relative position in the hierarchy being stable across time.

### The hierarchy of neural timescales at rest is dependent on behavioral demands

Next, we showed that neural timescales are task-dependent in general. Notably, we found that the baseline itself is not “intrinsic” but rather a reflection of the contextual cognitive and perceptual demands imposed on the brain. To test if resting-baseline neural timescales (i.e., neural timescales at rest) that are supposed to reflect a neuron and brain area’s innate temporal integration characteristics, are also reflective of the behavioral demand dependency described in the previous section, we estimated neural timescales during task-free periods of time. We did not impose any task demands on our animals, and there was hence no predefined resting-baseline before a trial (i.e., the intertrial interval was self-imposed). As a result, there was a wide repertoire of behaviors in the moments before a lever press (e.g., sitting, walking, etc.). To capture moments when the animal was at “rest”, we operationalized task-free trials as periods of 5 s (or longer) during which displacement of the 3D center-of-mass was less than 40 cm, excluding task engagement (see **Fig. 3A** and **Methods**). The average task-free period length was 6.5 s (SD = 0.4 s).

**Fig. 3.**
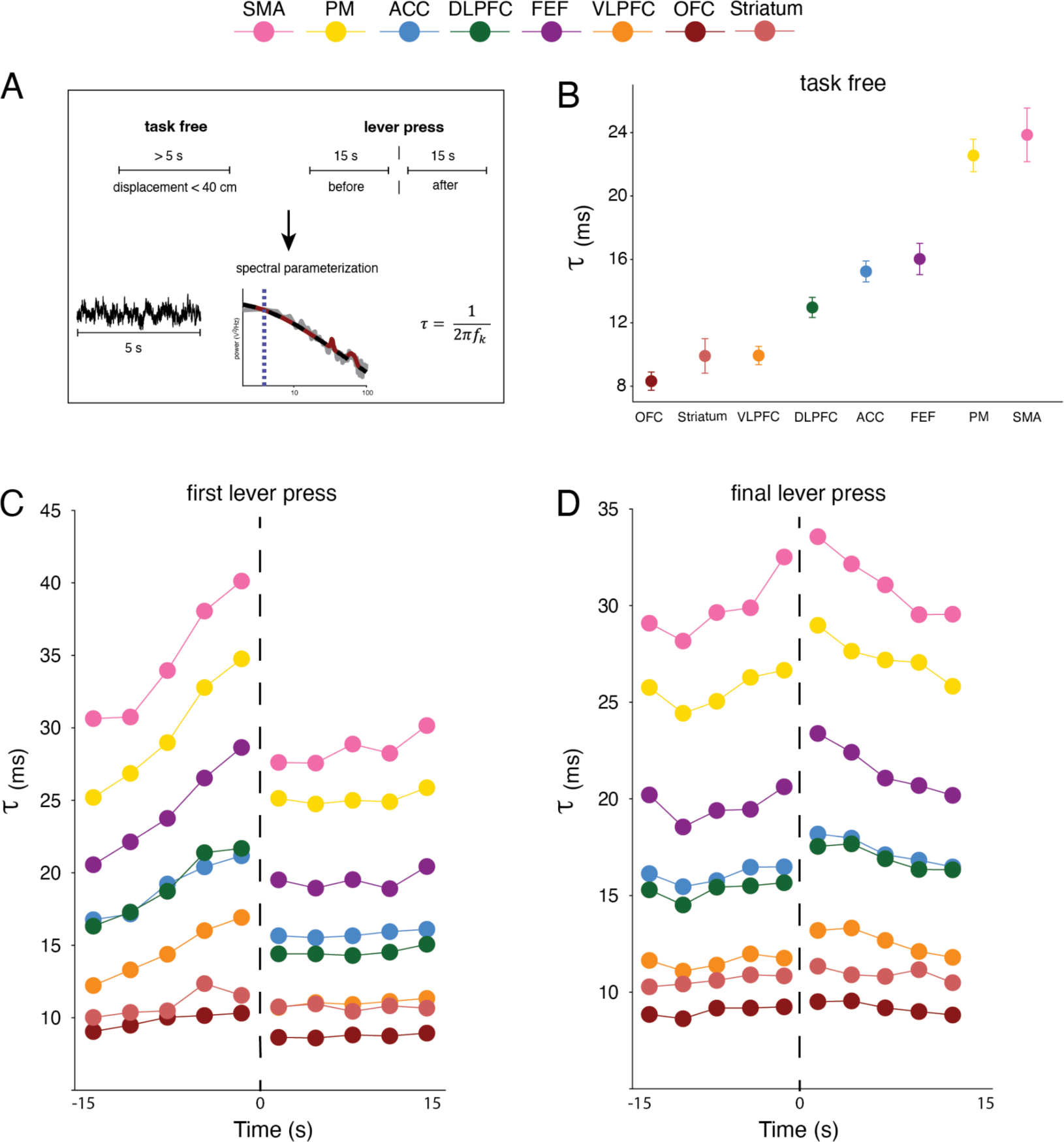
Neural timescales for events with different behavioral contexts. **(A)** Neural timescales (τ) estimation for task-free periods and lever presses. **(B)** Task-free neural timescales are hierarchically organized. **(C)** Neural timescales surrounding the first lever press. Vertical dotted line: time of lever press. **(D)** Neural timescales surrounding the final lever press. Vertical dotted line: time of lever press.

Using this approach, we showed that the hierarchy of task-free neural timescales was consistent with our session-wide results. We found the following hierarchy when the animal was at “rest”: OFC < Striatum - VLPFC < DLPFC < ACC < FEF < PM - SMA (**Fig. 3B**). The magnitude of the neural timescales differed significantly between areas, except for the following pairs: PM-SMA, ACC-FEF, VLPFC-striatum (p < 0.05, pairwise two-sided Mann-Whitney U-test computed across sessions with Bonferroni correction).

### Neural timescales changes are dependent on the behavioral context

Given the variability in neural timescales described above, we hypothesized that changes in neural timescales could depend on the behavioral context. In our experiment, the primary behavioral demands on the monkeys resulted from the pattern of engagement with the reward stations, i.e., the decisions to engage or disengage with a particular reward station location. We therefore divided events with lever presses into three categories: (1) first press on a feeder, (2) intermediate presses and (3) final lever presses. We hypothesized that these three types of lever presses could be associated with different neural timescales signatures because, even though they have identical actions, they have inherently different cognitive meanings and behavioral contextual demands. The first lever press reflects the decision to forage at a given reward station while the final lever press ends that goal sequence and requires an animal to decide on what to do next. Intermediate lever presses are more heterogeneous in terms of their position in the goal sequence and were therefore not considered for this analysis.

By estimating neural timescales before and after the first and last lever presses (**Fig. 3A**), we found an expansion in the magnitude of timescales during task engagement without an associated change in the relative position of an area in the overall hierarchy - i.e., all areas showed significant τ increase compared to resting-baseline (defined as the median task-free timescales at rest) at any time point before and after the event (p < 0.01, one-sided Wilcoxon signed-rank test with Bonferroni correction).

Interestingly, a clear and unique temporal dynamic profile separated the two categories of lever presses. For the first lever presses, we found that neural timescales gradually increased in the seconds leading up to the interaction with the reward station and sharply decreased in the seconds after (**Fig. 3C** and **Fig. 5B**). To assess the significance of this linear increase for individual areas, we conducted a linear regression model for each event with time as the predictor and neural timescales as the dependent variable. For all areas across events, the resulting regressions coefficients were significantly larger than 0 (p < 0.05, Wilcoxon signed rank test). After the first lever press, in some areas such as SMA, FEF, ACC, VLPFC and striatum neural timescales started increasing again while for the others, there was no significant trend across time (p < 0.05, Wilcoxon signed rank test). For the final lever presses, neural timescales gradually increased in the seconds leading up to and continued to increase after the interaction with the reward station (**Fig. 3D** and **Fig. 5C**). Similar to the first lever presses, we found that for all areas across events, the resulting regressions coefficients were significantly larger than 0 before the event (p < 0.05, Wilcoxon signed rank test). In contrast, in all areas we found that regression coefficients are significantly smaller than 0 after the event (p < 0.05, Wilcoxon signed rank test).

So far, we have therefore demonstrated a nested correspondence between neural timescales and the temporal scales over which behavior unfolds. At long (session-wide) temporal scales, neural timescales corresponded with overall task engagement as shown above, while at short temporal scales, neural timescales exhibited variability corresponding to ongoing behavioral demands from our task.

### The temporal adaptation of neural timescales varies by area

In the previous section, we demonstrated a correspondence between neural timescales and behavioral contextual demands. Although we showed that the hierarchy expands in the seconds leading up to the lever presses, we did not assess whether this magnitude change differed by area. Here, we hypothesized that this magnitude expansion of neural timescales might indeed be differentially modulated by differing behavioral demands per area. To that end, we examined the pairwise differences between areas (two-sided Mann-Whitney U-test) for the change from resting-baseline for each time point before and after the first and final lever presses (**Fig. 4**). Note that we chose this statistical approach for the sake of robustness and to avoid overfitting. We obtained the change in neural timescales per area by subtracting the respective area-specific resting-baseline (Δτ) for each time point. We found that this normalized adaptation differed in magnitude by area, such that while the change was undifferentiated several time points before the event, a differentiated hierarchy emerged just before the events of interest.

**Fig. 4.**
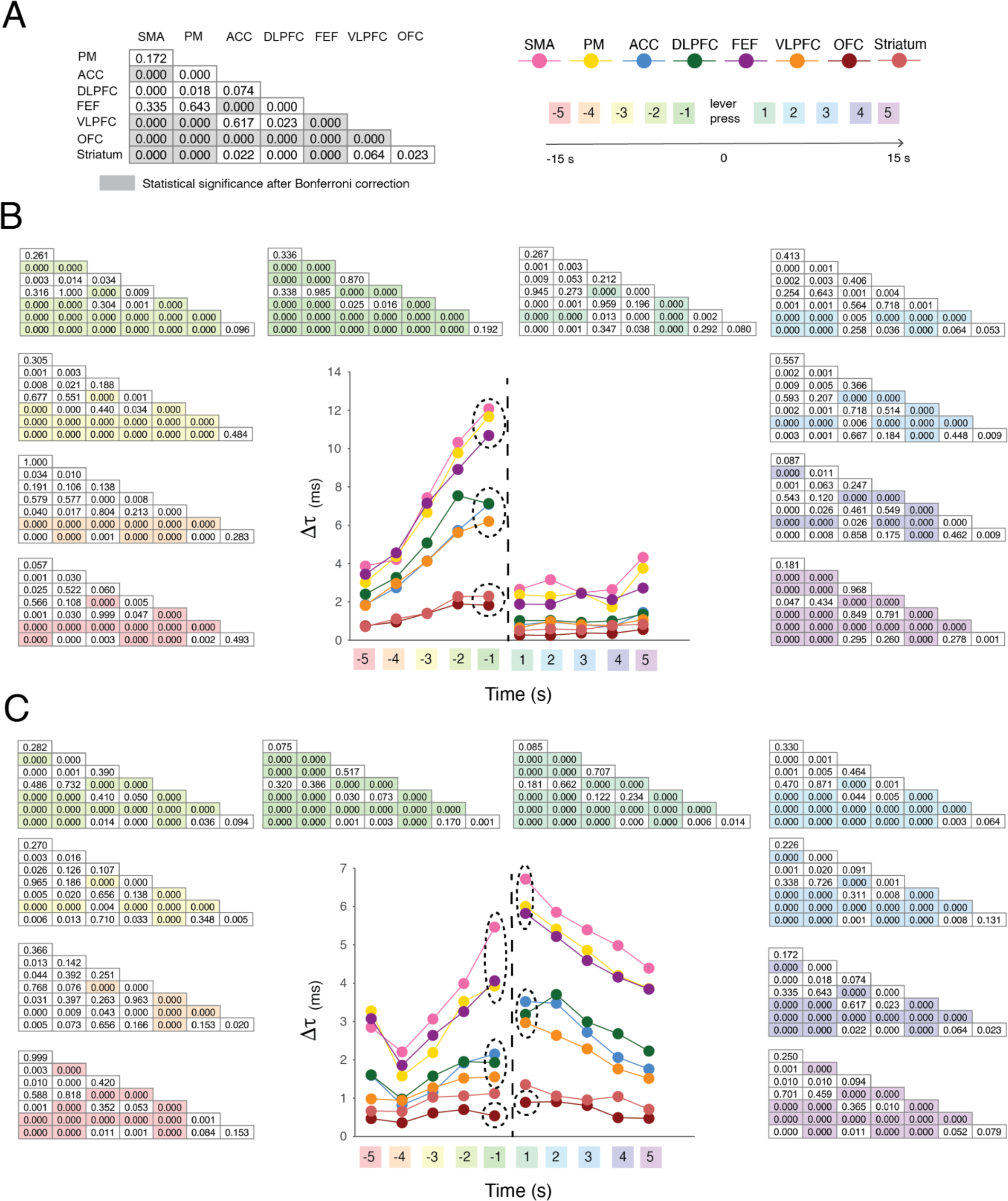
The normalized area-specific adaptation of neural timescales. (A) We examined pairwise differences between areas’ change from the area-specific resting-baseline (Δτ) for each time point before and after the event of interest. Left: example statistical table with the shaded areas representing statistical significance at p < 0.05 after Bonferroni correction. Right: color scheme of the individual areas and the color scheme of different time points (Note: used for indicating the associated statistical table). (B) The change from the area-specific resting-baseline for the time points before and after the first lever press. (C) The change from area-specific resting-baseline for the time points before and after the final lever press.

Notably, we found that a ventral to dorsal grouping of the areas emerged before each event: (1) the striatum and OFC displayed the smallest magnitude change in neural timescales; (2) the VLPFC, DLPFC and ACC displayed an intermediate level of magnitude change that was significantly higher than in the striatum and OFC; (3) and the FEF, PM and SMA displayed the significantly highest level of magnitude change —hence, these areas were placed at the top of the hierarchy of change (see table inserts in **Fig. 4** for statistics). We found this grouping before both first and final lever presses. However, each category of events exhibited unique temporal profiles after the event. Notably, for the first lever presses, the areas clustered right before the event and became undifferentiated immediately after (see table inserts in **Fig. 4** for statistics). For the final lever presses, similar clusters to those observed for the first lever presses emerged before the event, but they persisted after the event. In summary, while task engagement generally expands the magnitude of neural timescales, this expansion was differentially modulated by the behavioral task demands - i.e., it depended on the type of event and the brain area. Finally, we demonstrated that not only neural timescales, but also their area-specific magnitude changes were hierarchically organized along a ventral to dorsal axis.

### The dynamics of neural timescales reflect key foraging events

Finally, we looked into whether timescale adaptations differed between events with lever presses that have the same position in the sequence but are followed by a different outcome. This way we aimed to further dissect behavior with respect to its cognitive meaning and behavioral demands. To operationalize this, we looked at the fourth lever press which can indicate different behavioral motifs depending on the sequence of foraging bouts. For example, a monkey can leave a feeder after four lever presses without performing the fifth press to time out the system to for example go to the next feeder or engage in different behavior. From the perspective of an action, there is no difference between timing the system out versus choosing to disengage early. We thus hypothesized that neural timescales could exhibit differentiable temporal profiles also to these more intricate behavioral sequence differences. Practically, we compared neural timescales on the fourth lever press when the monkey decided to leave versus when they decided to stay for a fifth lever press (see **Fig. 5A**). Hence, we hypothesized that the “leave” before-after dynamics would be similar to that of the final lever press. In contrast, we hypothesized that there would be no significant before-after differences for the “stay” lever presses. For each event and area, we therefore computed the pairwise differences between before and after changes in the magnitude of neural timescales from resting-baseline. In other words, this analysis compared the timepoint proceeding and following a lever press event in terms of their magnitude change from resting-baseline. We found that for all areas, the before-after change from resting-baseline to the first lever press was characterized by a significant attenuation, with slower neural timescales before and faster neural timescales after the event (p < 0.05, one-sided Wilcoxon signed-rank test; **Fig. 5B**). The final lever press displayed the opposite pattern, with a significant increase in the change from resting-baseline, and ultimately slower neural timescales after the event (p < 0.05, one-sided Wilcoxon signed-rank test; **Fig. 5C**). As hypothesized, “stay” lever presses did not elicit significant changes in before-after dynamics - i.e., the event was not accompanied by a unique neural timescales signature (p < 0.05, two-sided Wilcoxon signed-rank test; **Fig. 5D**). In contrast, leave lever presses were accompanied by a significant before-after expansion of the magnitude of neural timescales which mimicked that of the final lever presses (p < 0.05, one-sided Wilcoxon signed-rank test; **Fig. 5E**).

**Fig. 5.**
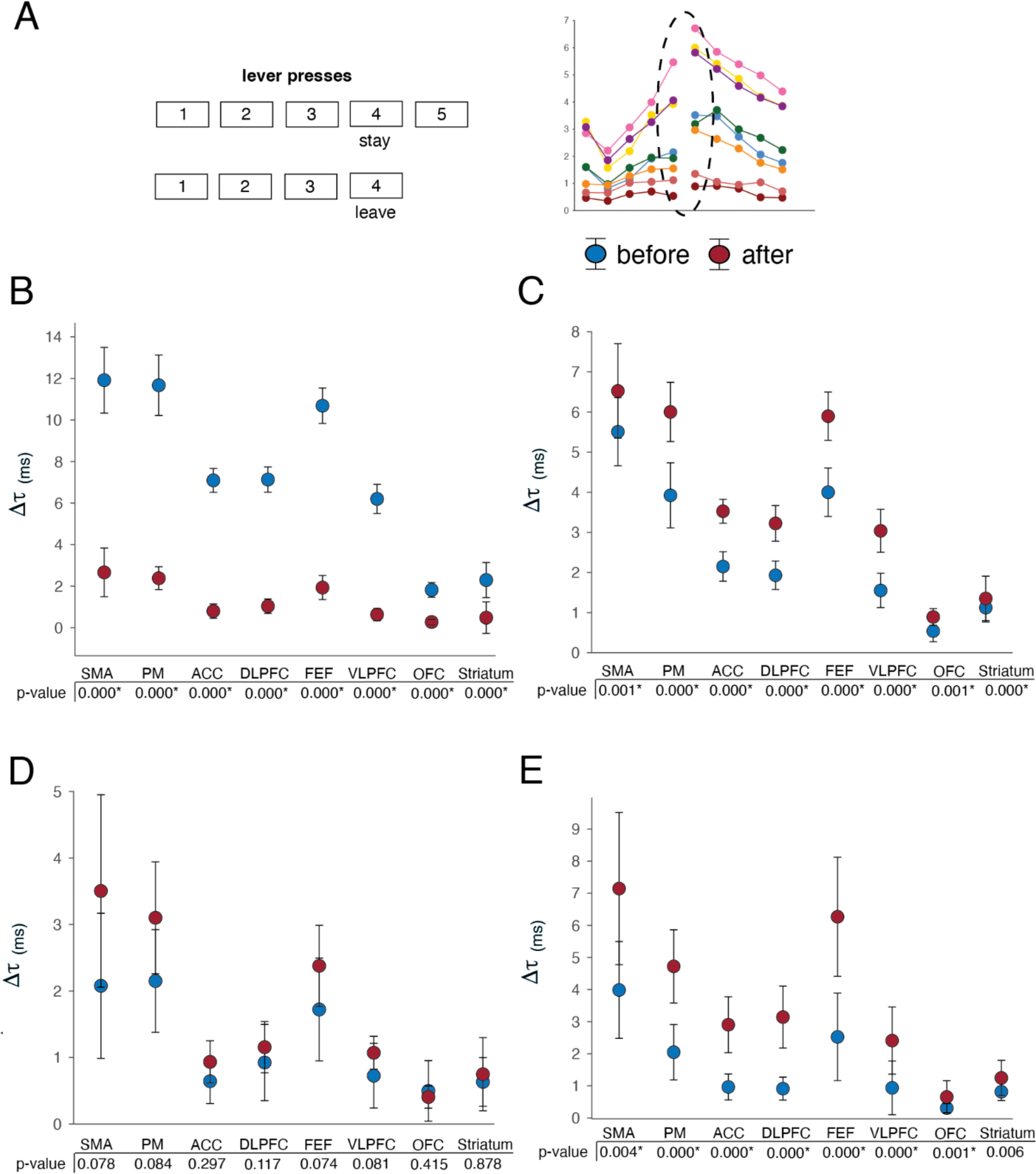
Neural timescales dynamics reflect fine grained abstract meaning. **(A)** Neural timescales (τ) estimation for three categories of lever presses: first lever press, intermediate (“stay”) lever press, final lever press. **(B)** The change in neural timescales from resting-baseline before and after the first lever presses. Asterisk: statistical significance at p < 0.05 after Bonferroni correction. Y-axis: change in neural timescales from resting-baseline (Δτ). **(C)** The change in neural timescales from resting-baseline before and after the final lever presses. Asterisk: statistical significance at p < 0.05 after Bonferroni correction. Y-axis: change in neural timescales from resting-baseline (Δτ). **(D)** The change in neural timescales from resting-baseline before and after the stay lever presses. Asterisk: statistical significance at p < 0.05 after Bonferroni correction. Y-axis: change in neural timescales from resting-baseline (Δτ). **(E)** The change in neural timescales from resting-baseline before and after the leave lever presses. Asterisk: statistical significance at p < 0.05 after Bonferroni correction. Y-axis: change in neural timescales from resting-baseline (Δτ).

## Discussion

The brain is characterized by a hierarchical gradient of neural timescales in both human and nonhuman primates (Ito et al., 2020; Manea et al., 2022; Raut et al., 2020), which is assumed to arise from macroscale and microcircuit anatomical and functional connectivity, as well as variation in cytoarchitecture (Murray et al., 2014; Chaudhuri et al., 2015). The question arises how these temporal properties of the brain give rise to adaptive behavior that requires flexible adjustment of temporal coding and integration demands. Here we found a ventro-dorsal hierarchy of neural timescales that is unique to this foraging environment. Importantly, we showed that this hierarchy is preserved even in the context of flexible task demands. However, the magnitude of neural timescales dynamically expanded depending on overall task engagement over long temporal scales, but also varied with the cognitive demands of our task over shorter temporal scales. Notably, we observed systematic changes in the magnitude of neural timescales that span the duration of a recording session —in both animals and during every recording session, both the magnitude of neural timescales and task engagement gradually and reliably decreased over time. Importantly, these results were not driven by variability in motor-related activity. Within these global session-wide changes, we found variability in the magnitude of neural timescales that is associated with the abstract cognitive meaning of the different foraging events. Hence, the neural timescales change patterns differentiated between fine-grained behavioral states. Our results are evidence that the multitude of external temporal scales over which behavior unfolds is mirrored by changes in neural timescales that occur at multiple scales, with local foraging-related changes nested within general engagement-related changes that span longer temporal scales.

Contrary to previous work on timescales, which was usually done in the context of memory-related or value-encoding tasks (for an overview, see Cavanagh et al., 2020; Wolff et al., 2022), we examined self-paced unconstrained behavior that was not focused on an isolated cognitive component. We consistently found a stable ventral to dorsal hierarchy of neural timescales that extended from OFC to motor-related areas. This was the case across all our analyses - i.e., for neural timescales estimated during unconstrained movement, around foraging events and even in the absence of task engagement. Our observed hierarchy partially overlapped but also deviated from previous work (see Fig. 1A for an overview of previous findings). The first major discrepancy was that the ACC, usually displaying the slowest timescales (Maisson et al., 2021; Murray et al., 2014), was not at the top of our observed hierarchy, but rather exhibited intermediate timescales. Moreover, motor–related areas, which usually display fast timescales in nonhuman primates (Cirillo et al., 2018), exhibited the slowest temporal dynamics in our foraging environment. This was actually in full agreement with results from a rodent paradigm that allows for whole-body movement (Pinto et al., 2022). We believe these results originate from our experimental paradigm that allowed for unconstrained movement in a large open space. This contrasted with studies using chaired paradigms that require head fixation and restrained body movement. In line with our initial hypothesis, our results supported the idea that environmental demands shape the observed hierarchy of neural timescales. This is not surprising since behaviorally-relevant neural timescales imply a certain amount of dynamic range. It is important to note that our results are not incompatible with the previous literature on hierarchies of neural timescales at rest but are rather complementary by investigating neural timescales in a new context, that of unconstrained behavior. Our results further expanded previous work as we simultaneously estimated neural timescales in the dorsal striatum and prefrontal cortical structures, giving us the opportunity to directly place the dorsal striatum in the context of the broader extensively studied cortical hierarchy. The striatal neural timescales reported here place the striatum on a comparable level to ventral prefrontal areas, which is in agreement with previous reports of neural timescales in this subcortical structure (Nougaret et al., 2021). Overall, we confirmed our hypothesis that the observed hierarchy of neural timescales emerges from the particular input-output demands imposed on the brain within the bounds of what the anatomical network permits.

Matching previous work on neural timescales at rest, i.e., estimated during the baseline or pre-trial period (for an overview, see Fig. 1A), we estimated neural timescales during immobility periods. Since the animals were free to decide when or even if, to engage in the task, the current paradigm did not have a traditional pre-trial period comparable to previous work —i.e., the repertoire of behaviors before engaging with a reward station is highly heterogeneous. Immobility periods, when the animal is disengaged from the task, were the closest match to capturing the animal at rest. It is important to note that rest or baseline is generally difficult to define —the traditional fixation period used to define neural timescales at rest is assumed to include little task-relevant signals. However, lack of outward behavior does not imply a lack of cognitive processing. Indeed, previous findings on the relationship between neural timescales at rest, estimated during the intertrial period, and strength of neural encoding during a task did not always replicate (Spitmaan et al., 2020). This might be a result of task-relevant cortical activity emerging or remaining during the fixation period. To estimate neural timescales at “rest”, work in anesthetized subjects or during sleep might be a viable alternative (for example, see Cushnie et al., 2023; Manea et al., 2022; Zilio et al., 2021), although these approaches come with their own disadvantages and confounds. We have previously shown that neural timescales estimated from fMRI data in anesthetized nonhuman primates replicated hierarchies derived from neuronal spiking data, although they did not perfectly match (Manea et al., 2022). Amongst many potential reasons for the observed deviations, one could be that anesthesia provides a special controlled state. Alternatively, electrophysiological recordings during dedicated rest periods, in the absence of any task, similar to the human resting-state fMRI literature, could shed light on this issue. We speculate that while anatomy constrains the space of possible neural timescales, contextual behavioral demands modulate the observed hierarchy even at rest. Interestingly, the hierarchy of neural timescales we observed mirrored functional hierarchies that we found in this dataset with respect to action (see Voloh et al., 2022) and spatial navigation encoding (see Maisson et al., 2022). In these studies, encoding of spatial navigation and action-related variables was progressively stronger from ventral to dorsal areas (Maisson et al., 2022; Voloh et al., 2022), and hence potentially facilitated by longer temporal processing or integration windows as reflected by slower neural timescales. Our current timescale findings and the previous task variable encoding findings, match our previous work demonstrating that the hierarchy of neural timescales at rest in the medial PFC closely follows a ventro-dorsal functional hierarchy of the decodability of choice-relevant task variables - i.e., encoding of task related variables is stronger in areas with longer neural timescales (Maisson et al., 2021).

We found that neural timescales expanded during task engagement, in agreement with previous studies (Gao et al., 2020; Zeraati et al., 2021). Interestingly, we found a monotonic relationship between the extent of task engagement and the overall magnitude of neural timescales over a long temporal scale which spans the recording session. More task engagement was accompanied by an expanded hierarchy of overall slower neural timescales throughout the recording session. Within this session-wide change with behavioral engagement, we demonstrated local event-related changes in the magnitude of neural timescales. Although the action of pressing a lever was similar irrespective of its location in the broader foraging context, unique temporal dynamics of neural timescales were associated with differences in cognitive meaning. We speculate that this is a result of differences in the underlying computations associated with these lever presses. Neural timescales thus seemed to track the temporal persistence of information relevant during the ongoing decision process in a behaviorally-relevant manner. For example, while the first lever press reflects the decision to forage at a particular reward station, the action per se can be seen as the end goal. Interestingly, this is accompanied by a significant drop in the magnitude of neural timescales. We speculate that accomplishing this goal could act as a stop signal for integration. In contrast, the last lever press seemed to reflect ongoing integration related to the animal needing to decide what to do next, with neural timescales continuing to expand after the event. Nevertheless, the specific meaning of these time-locked local changes in the magnitude of neural timescales remains an outstanding question. Although ecologically valid, unconstrained behavior automatically introduces variability that cannot be controlled for, nor is easily modeled. Our results therefore could be extended by directly manipulating the information integration across multiple temporal scales, while introducing clear task endpoints.

Despite widespread modulation by general task demands, we also found that the temporal dynamics of neural timescales showed time-locked changes to certain foraging events that differed by area. We found a significant differentiation across areas in the time-locked expansion of hierarchies linked to certain foraging events. Notably, however, the ventral to dorsal grouping of areas was preserved during this expansion - i.e., the dorsal striatum and OFC exhibited the lowest level of change from resting-baseline; VLPFC, DLPFC and ACC, exhibited intermediate levels; and FEF, PM and SMA were at the top of the hierarchy, displaying the highest level of change from resting-baseline. Importantly, although this ventro-dorsal hierarchy of change was present for all lever presses, its emergence and persistence depended on the cognitive meaning mentioned above. Changes in the real world happen on different timescales and these results suggest that individual brain areas may contribute distinctly to behavioral adjustments on different temporal scales. It is not trivial to quantify unconstrained behavior, and it is even more challenging to infer the cognitive state of the animal in this setting. As a result, it remains an open question as to what particular task variables are associated with the observed variability.

Here we use LFP rather than single-unit activity to infer timescales for two reasons. First, LFP activity, in our dataset, offered much broader spatial coverage and provided us with the ability to record a large number of areas simultaneously. That is, we were able to leverage the nature of this signal to infer neural timescales in all areas across time throughout the recording session. Second, the neuronal firing rates in this dataset are sparse and hence, it would have been difficult if not impossible to calculate the autocorrelation function across time for many cells. Although LFP and single-unit activity are fundamentally different signals, it has been shown that they have related neural timescales at rest, and that they are both hierarchically organized in the nonhuman primate cortex (Gao et al., 2020). This is also the case for neural timescales estimated from neuronal spiking and fMRI data (Manea et al., 2022). Nevertheless, the exact correspondence between neuronal spiking, LFP and BOLD timescales and the extent of their overlap remains unclear.

In conclusion, we demonstrated that neural timescales vary with different indices of task engagement that are an aggregate of behavioral state parameters broadly encompassing foraging-related behaviors, action and spatial navigation encoding. Despite the magnitude of neural timescales expanding with task demands and engagement, we found a stable hierarchical ordering of the areas’ neural timescales. We not only provided evidence for the context-dependence of neural timescales, but we also demonstrated that these temporal dynamics were complex and behaviorally relevant. Further investigations and careful experimentation that manipulate the temporal scales over which an animal has to integrate information are needed to better understand the link between neural timescales and the temporal scales over which behavior unfolds.

## Methods

### Surgical procedures

Animal procedures were designed and conducted in compliance with the Public Health Service’s Guide for the Care and Use of Animals and approved by the Institutional Animal Care and Use Committee of the University of Minnesota. Two male rhesus macaques (Macaca mulatta) served as subjects. Animals were habituated to laboratory conditions, trained to enter and exit the open arena, and then trained to operate the water dispensers. We placed a cranial form-fitted Gray Matter (Gray Matter Research) recording chamber and a 128-channel microdrive recording system (SpikeGadgets) over the area of interest. We verified positioning by reconciling preoperative MRI as well as naive skull computed tomography images (CT) with postoperative CTs. Animals received appropriate analgesics and antibiotics after all procedures. The planning of the chamber and subsequent image alignment was performed in 3D slicer. Brain area segmentation followed the macaque D99 parcellation in NMT space (Saleem et al., 2021)

### Electrophysiological recordings

We recorded with a 128-channel microdrive system (Gray Matter Research), targeting a wide swath of the prefrontal cortex ranging from OFC to PM, and the striatum. Each electrode was independently moveable along the depth dimension. Neural recordings were acquired with a wireless data logger (HH128; SpikeGadgets). The data logger was triggered to start recording with a wireless RF transceiver and periodically received synchronization pulses. Data were recorded at 30 kHz, stored on a memory card for the duration of the experiment, and then offloaded after completion of the session. Each reward station had local code running the experiment. Task events triggered a TTL pulse, as well as a wireless event code. A dedicated PC running custom code controlled all reward stations, and aggregated event codes. Syncing of all data sources was accomplished via the Main Control Unit (MCU; Spikegadgets), which received dedicated inputs from the pose acquisition system (see below), and reward stations. Recording sessions were initiated and controlled by Trodes software (Spikegadgets). After neural recordings were offloaded, they were synced with other sources of data via the DataLogger GUI (Spikegadgets).

We recorded for 4-6 days weekly for a period of 4-6 months. For an initial period of 2-4 weeks, we lowered up to 10 electrodes in each session until each had punctured the dura and their position was well-within cortex as confirmed from the MRI reconstruction. Subjects still performed experiments, but as the signal was noisy, no recordings were performed during this time. A typical recording day consisted of multiple stages, including electrode adjustment, an experimental session, and extraction of the recorded signal. For the duration of the experiment, on each day, we tracked yields on each electrode and visually assessed the quality of the signal. If an electrode had poor yields for up to 5 days in a row, we would lower it up to 1 mm (or more if it was intended to move to a new area).

To obtain local field potentials (LFP), we bandpass filtered the raw signal with a second order, two-pass Butterworth filter and Hand taper in the range [0.1 300] Hz. Only recordings that showed evidence of neural unit activity (confirmed with separate modified spike sorting analysis) were used for further analysis. Subdivisions of the brain were collapsed to anatomical areas, listed below as defined in the D99 parcellation of the NMT atlas (Saleem et al., 2021): ACC: 24a’, 24a, 24b, 24b’, 24c, 24c’; VLPFC: 45a, 45b, 46d, 46v, 46f, 12r; DLPFC: 8bd, 8bs, 9d, 8bm, 9m; FEF: 8ad, 8av; SMA: F3, F6; PM: F1, F2, F5, F7, F4; OFC: 13b, 13m, 13l, 12l, 12m, 12o, 11l, 11m; Striatum.

### Behavioral tracking

We developed a system that can perform detailed three-dimensional behavioral tracking in rhesus macaques with high spatial and temporal precision (Bala et al., 2020). The system uses 62 cameras positioned around a specially designed open field environment (2.45 × 2.45 × 2.75 m) with barrels (4 barrels located in the corners; height: 78.8 cm; diameter 46.5 cm) in which macaque subjects can move freely in three dimensions and interact with computerized reward stations.

### Pose acquisition and reconstruction

A detailed protocol of the pose acquisition, reconstruction preprocessing can be found in Voloh et al. (2022) and Maisson et al., (2022).

### Behavioral task

The environment contained four reward stations (“patches”) that dispensed water with a programmed delivery schedule. The reward stations were rectangular white boxes with a display monitor placed in the middle, a lever to the left and a waterspout to the right. The display monitor indicated the availability of the station for foraging (solid blue), reward delivery (solid white background with a solid green cross) or unavailability of the station (i.e., the timeout period; solid white). Each station delivered a fixed amount of water (1.5 ml) per lever press. At any given time, each of the first four lever presses were rewarded and the fifth lever press led to a 3-minute timeout period (i.e., depleted station). The subjects could freely decide when and how to interact with the reward stations. No reset or deactivation was applied if the animal left the patch. The timeout could only be triggered after four rewarded and one unrewarded lever press. Each rewarded lever press followed the same programmed sequence. The availability of the reward station was indicated by a solid blue display. A lever press changed the display to white with a green cross in the center, the auditory cue was played, and the solenoid opened to dispense reward. After dispensing, the solenoid closed, the auditory cue ended, and the green cross disappeared. The display remained white for two additional seconds before it turned blue again. The fifth lever press was instead followed by the screen immediately turning white, with no visual or auditory reward cue and no reward delivery. Other than the interaction with the reward stations, the measured behavior was the subject’s unconstrained movement.

### Behavioral variables

#### Speed of movement

For this analysis we used the 3D center-of-mass (defined as the midpoint between the hip and neck joint) trajectories. We calculated speed as the magnitude of the numerical derivative of the 3D center-of-mass of the subject.

#### Task-free trials

Task-free trials were operationalized as windows longer than 5 seconds when the animal was relatively immobile (i.e., the 3D center-of-mass displacement was less than 40 cm). Task engagement (i.e., continuous interaction with a reward station) was excluded.

#### Lever presses

We divided lever presses into three categories: first, intermediate and final lever presses. The first lever press was operationalized as the first interaction with a reward station after the animal changed stations. The final lever press was operationalized as any interaction with a reward station before changing stations. We took advantage of the fact that at times subjects prematurely abandoned the reward station after the fourth lever press. Specifically, we compared responses on the fourth lever press when the subject decided to leave versus when they decided to stay for a fifth lever press. The stay lever presses fit within the intermediate category.

### Timescales estimation PSD

PSDs were estimated using the conventional Welch’s method, where short-time windowed Fourier transforms are computed from time series and the mean is taken across time. We used 1 s long Hamming windows with 500ms overlap.

### Spectral parametrization

Spectral parameterization (Donoghue et al., 2020)was applied to extract timescales from PSDs. Briefly, the log-power spectra were decomposed into a summation of narrowband periodic (modeled as Gaussians) and aperiodic (modeled as a Lorentzian function centered at 0 Hz) components. To infer timescales, the periodic components are discarded, and timescales were inferred from the aperiodic component of the PSD. Specifically, τ can be estimated from the parameter *k* as 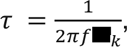 where 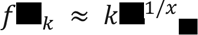 is approximated to be the knee frequency, at which a “knee” in the power spectrum occurs. For a detailed mathematical description of the model and the timescale inference technique, see Donoghue et al., 2020 and Gao et al., 2020.

### Neural timescales

Timescales were inferred for each channel individually and then collapsed across channels within an area by taking the median in order to limit the impact of outliers. We excluded windows for which the PSD parameterization failed or for which the model fit (i.e., R squared) was lower than 0.8. Channels with a failure rate higher than 20% were excluded from further analyses.

#### Session-wide

Timescales were inferred by applying spectral parameterization (see above) to the entire session, using a 10 s moving window with 5 s overlap.

#### Event-related

Timescales were inferred 15 s before and after the interaction with a reward station. We inferred timescales by using a 5 s moving window with 2.5 s overlap. We excluded windows for which the PSD parameterization failed or for which the model fit (i.e., *R*^2^) was lower than 0.8.

#### Task-free

Timescales were estimated across time over the length of the task-free windows by using a 5 s moving window with 2.5 s overlap over the length of the trial. We excluded windows for which the PSD parameterization failed or for which the model fit (i.e., *R*^2^) was lower than 0.8. To obtain one value per trial, we subsequently averaged the neural timescales estimated for any given task-free trial. For each session, the median task-free neural timescales at rest were used as the baseline for further analyses.

### Statistical analysis

We used Pearson correlation coefficient to estimate the relationship between number of lever presses, speed and neural timescales across individual recording sessions.

Given that neural timescales were not normally distributed, we opted for nonparametric tests to assess statistical significance. We used Wilcoxon signed-rank tests to assess the statistical significance within individual areas - i.e., the change from resting-baseline and the change in neural timescales between neighboring time points. We used Mann-Whitney U-test to assess the statistical significance between brain areas - i.e., the difference in resting-baseline (or task-free timescales) and the difference in change from resting-baseline around the events of interest. To correct for multiple comparisons, we applied Bonferroni correction.

### Multiple regression analysis and subsampling

To perform this analysis, we only included sessions for which the lever press events, behavioral tracking, and neural timescales were available. The total number of data points per area can be calculated as the number of sessions x number of segments (i.e., 10). To quantify the effect of task engagement (i.e., the number of lever presses in any given segment) and movement of speed on neural timescales, we used a subsampling procedure to estimate the average effect and the confidence interval of these predictors. For each area, we randomly sampled without replacement n (i.e., equivalent to the number of sessions) observations out of the total number of data points. For each subsample, a linear regression model was fitted, with the number of lever presses and speed as regressors, and neural timescales as the response variable. For each area, we repeated the procedure 1000 times. For each predictor, we assessed the difference between areas using an independent sample t-test. For each area, we assessed the difference between predictors using a paired t-test. To correct for multiple comparisons, we applied Bonferroni correction.

### Bayesian regression model

To assess the relationship between brain areas and session-wide neural timescales, we modeled the predictor (i.e., brain area) as a monotonic effect (Bürkner & Charpentier, 2020). This approach is advantageous for ordinal predictors, in this case the hierarchical organization of brain areas, without falsely treating them as continuous, unordered categorical variables or ordered categorical variables with equidistant levels. In this approach, one estimates one parameter (*b*) which captures the direction and size of the effect - i.e., average increase/decrease in the dependent variable associated with the variable. Additionally, one estimates the percentages of the overall increase/decrease that is associated with each of the differences between neighboring variable levels - and hence, these parameters determine the shape of the monotonic effect. For a more detailed explanation, see Bürkner et al., 2020. Brain area was modeled as a monotonic effect and session-wide neural timescales served as the dependent variable.

## Acknowledgements

We thank the Hayden/Zimmermann lab for valuable discussions as well as Brenna Knaebe for help with animal care and preparation. This work was supported by NIH grants R01 MH128177 (JZ), P30 DA048742 (JZ, BH, AZ), R01 MH125377 (BH), NSF 2024581 (JZ, BH) and a UMN AIRP award (JZ, BH, AZ) from the Digital Technologies Initiative (JZ), from the Minnesota Institute of Robotics (JZ).

---

**Fig. 2 Supplement 1.**
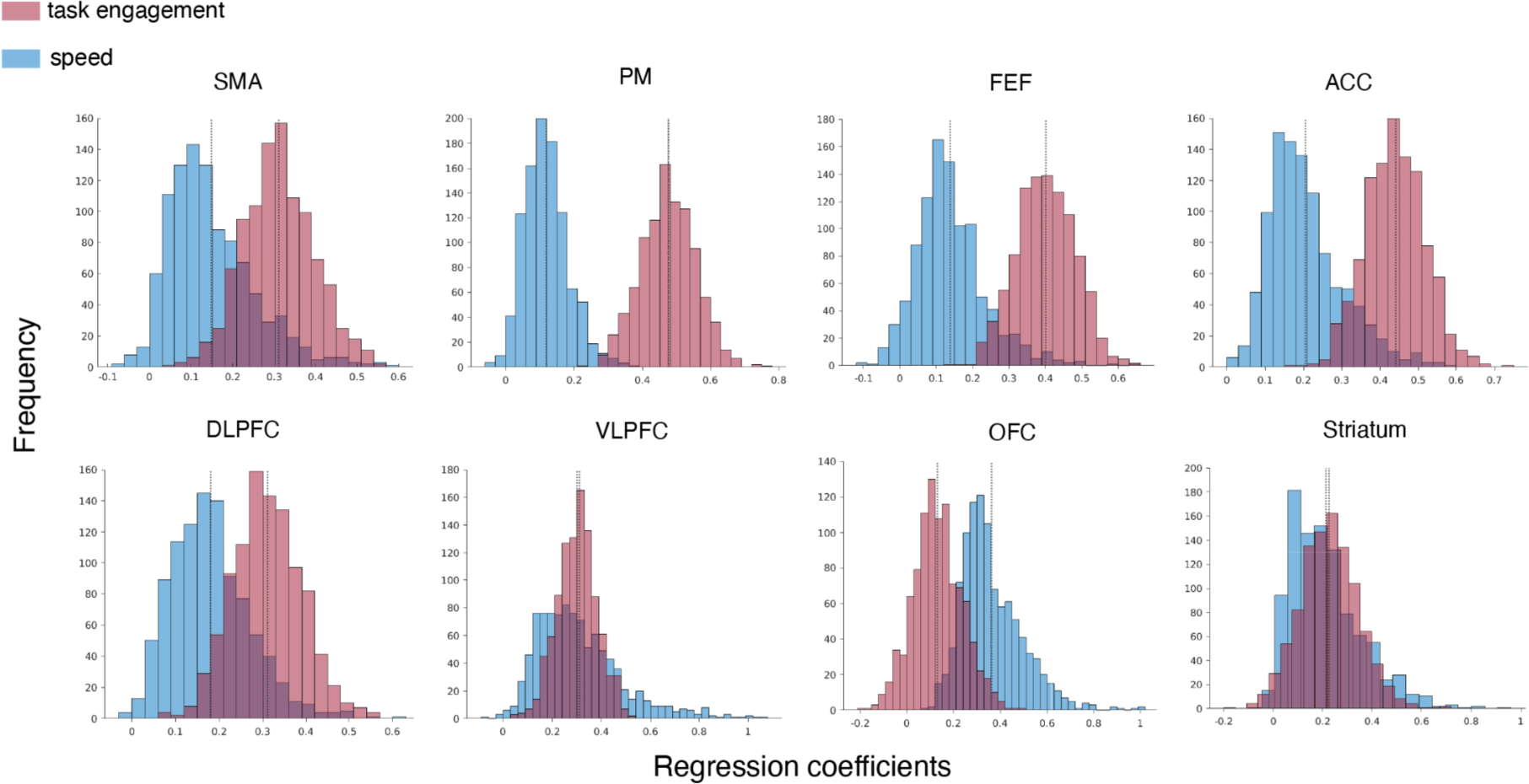
The distribution of standardized β coefficients for task engagement and speed

